# Grey and White Matter Pathways Underlying the Relationship Between Aerobically Active Lifestyle, Cardiorespiratory Function, and Episodic Memory in the Aging Human Brain

**DOI:** 10.1101/2020.12.20.423704

**Authors:** Tamir Eisenstein, Nir Giladi, Ofer Havakuk, Yulia Lerner

## Abstract

Aging is associated with structural alterations of the hippocampus, a key region in episodic memory processes. Aerobic activity and maximal aerobic capacity (MAC), a key measure of cardiorespiratory function and a physiological adaptation of aerobic exercise, have been associated with biological and cognitive resilience of the brain. However, investigations of their relationship with the hippocampus in humans had resulted with inconsistent findings. This study aimed to investigate the relationship between lifestyle’s aerobic activity and MAC and hippocampal grey and white matter structure, as well as episodic memory performance in cognitively healthy older adults. In addition, we examined the relationship between aerobic activity and MAC, and cerebrovascular pathology expressed as white matter lesions (WML). Next, we used a regression-based mediation analysis to examine possible biological pathways which may underlie the relationship between MAC and hippocampal volume, which was demonstrated in previous works, and was confirmed in the current study. Fifty cognitively healthy older adults (70.92 ± 3.9 years) were divided into aerobically active (n=27) and non-active (n=23) groups, and performed structural and diffusion MRI. Forty-two participants were also evaluated for MAC. Aerobically active lifestyle and higher MAC were associated with increased hippocampal volume and microstructural integrity, as well as increased fornix microstructural integrity, and lower WML burden (*p*<.05). In addition, both factors were correlated with increased episodic memory performance (*p*<.05). Mediation analysis revealed two pathways potentially mediating the relationship between MAC, hippocampal volume, and episodic memory – a white matter pathway consisted of WML and fornix microstructure, and grey matter pathway including hippocampal microstructure. These findings shed light on possible neurobiological mechanisms that could potentially underlie the neuroprotective effect of cardiorespiratory function and aerobic physical activity on hippocampal macrostructure and memory function in the aging human brain.

## 1. Introduction

Aging has been demonstrated to be accompanied by morphological alterations in the human brain. Magnetic resonance imaging (MRI) studies have demonstrated progressive structural changes of neural tissue expressed both at the macro- and microstructural levels. Global and regional decreased volumes of grey and white matter have been demonstrated with increased age ^1,2^, while neuroimaging markers derived from diffusion MRI that reflect the microstructural integrity of the tissue such as fractional anisotropy (FA) and mean diffusivity (MD) have also been shown to be affected during aging ^3^. While FA reflects the directionality of the diffusion of water molecules, and mostly implemented in measuring the integrity of white matter fiber tracts, MD reflects the average degree of diffusion, and may indicate microstructural changes in the grey matter ^4^. Since breakdown of cellular barriers to diffusion (e.g., cellular membranes) may underlie increased MD in the tissue grey matter, this measure could provide valuable information about cellular damage and the microstructural integrity of grey matter ^4,5^.

One region that has been demonstrated to exhibit significant structural changes during aging is the hippocampus, which plays a key role in episodic memory processes. Decreased hippocampal volume has been repetitively documented in older adults ^1,2,6,7^, and hippocampal grey matter MD has been shown to increase with age ^8^, possibly as early as the fifth decade to life ^9^, correlating with multidimensional episodic memory performance. Moreover, in comparing to hippocampal volume, hippocampal MD has been demonstrated to be a more sensitive biomarker of progression to Alzheimer’s disease dementia, and to episodic memory deficits in patients with amnestic mild cognitive impairment ^10^, as well as in cognitively-intact older adults ^11^. In addition, morphological changes in hippocampal-related white matter areas have also been shown in older adults including decreased volume and lower FA values in parahippocampal white matter ^12,13^ and structural alterations in the fornix ^14,15^. A potential mechanism which may underlie, at least in part, the progressive deterioration observed in hippocampal structure during aging may originate in accumulating cerebrovascular pathology. White matter lesions (WML) are an MRI biomarker of cerebrovascular insult and cerebral small vessel disease visible as hypo-intense white matter foci on T1-weighted images or hyper-intense white matter areas on T2-weighted images ^16^. WML burden has been associated with increasing age ^17,18^, cognitive decline and dementia ^19^. However, its relationship with hippocampal structure is still not clear, and while some works demonstrated an adverse association between WML and hippocampal volume ^20–22^, others found no connection ^23,24^. Recently, WML load have been shown to correlate with decreased hippocampal volume and atrophy rate in cognitively-intact older adults, while hippocampal volume was also found to be more sensitive to WML load than whole brain volume ^25^.

Several lifestyle factors have been suggested to be associated with neurobiological and cognitive resilience in face of age-related deterioration, one of which being physically active lifestyle ^26–28^. However, studies examining the effect of aerobic exercise on the hippocampus in humans demonstrated inconsistent results ^29^. While, several human studies did demonstrate positive association between aerobic exercise and hippocampal structural integrity following exercise intervention ^30–32^, others have found no specific effect of aerobic exercise ^33,34^. Interestingly, in studies which did found positive associations, the extent of these improvements was shown to be correlated with cardiorespiratory physiological adaptations, primarily increased maximal oxygen consumption. This physiological parameter expresses the highest metabolic rate of the body to generate adenosine triphosphate (ATP) molecules through aerobic metabolism ^35^, or maximal aerobic capacity (MAC). MAC has also been demonstrated to be associated with cerebrovascular integrity in older adults, and linked to hemodynamic functions such as vaso-reactivity and cerebral blood flow velocity ^36,37^. In addition, while individual studies demonstrated inconsistent results regarding the relationship between MAC and WML burden, a small but significant negative relationship was detected between higher levels of aerobic exercise/MAC and decreased WML burden in a previously performed meta-analysis ^38^, suggesting for a potential moderating effect of aerobic activity and MAC on cerebrovascular pathology.

As the importance of physically active lifestyle in promoting neuroprotection and neurocognitive resilience in aging is increasingly recognized, the neurobiological mechanisms underlying these effects are still a matter of delineation. Furthermore, while several potential mechanisms have been suggested to mediate exercise-dependent effects on hippocampus and memory based on animal models ^39^, human research generally lacks the ability to directly explore potential microscale biological mechanisms on the molecular or cellular level. Nonetheless, the current diversity of MRI modalities enables us to examine several levels of neurobiological measures and base our mediation investigation on designated statistical methods. Hence, the aims of the present study are several-fold: first, examining and confirming a neuroprotective relationship between aerobically active lifestyle and hippocampal grey and white matter structures, cerebrovascular pathology, and episodic memory; second, as MAC has been suggested to mediate (at least partially) the neurocognitive effects of aerobic exercise, we aimed to investigate the dose-response relationship between MAC and all aforementioned neurocognitive indices; third, we aimed to investigate whether cerebrovascular pathology and hippocampal grey and white matter microstructure may mediate the relationship between MAC and hippocampal macrostructure (volume); and last, we aimed to identify biological pathways that may underlie and mediate the relationship between MAC and episodic memory performance.

## 2. Materials and methods

### 2.1 Participants

Fifty older adults aged 65-80 (mean age 70.92 ± 3.9 years) were recruited for the study (22 women/28 men). All participants were recruited from the community, were fluent Hebrew speakers, and reported no current or previous neuropsychiatric disorders (including subjective cognitive decline or depressive symptoms) or any other current significant unbalanced medical illness (e.g., cardiovascular or metabolic). The research was approved by the Human Studies Committee of Tel Aviv Sourasky Medical Center (TASMC), and all participants provided written informed consent to participate in the study. Each participant arrived at the lab on two separate days, no longer than 3 weeks apart. Participants were asked not to change their physical activity habits between the two sessions and to report any unexpected medical events during this period. On the first meeting, a neuroimaging session was conducted, while on the second meeting, a memory assessment was first conducted followed by a cardiopulmonary exercise test to prevent any exercise effects on cognitive performance.

### 2.2 Aerobic activity lifestyle assessment

Current aerobically active lifestyle was assessed using a background interview that examined lifestyle activities, and specifically assessed the aerobic exercise habits of the participants during the last year. Next, participants were divided into two groups, active (n=27) and non-active (n=23) based on their reported activity patterns. To avoid potential information bias from participants’ reports on active lifestyle, the active group consisted of participants who declared performing routinely aerobic activity (e.g., walking, jogging, running, cycling, dancing) at least twice a week during the last year.

### 2.3 Maximal aerobic capacity assessment

Forty-two participants (16 women/26 men) also underwent a graded maximal cardiopulmonary exercise test performed on a cycle ergometer to evaluate MAC. Assessments were conducted at the Non-Invasive Cardiology Outpatient Clinic at TASMC. Tests were supervised by a cardiologist and an exercise physiologist while continuously monitoring for cardiopulmonary parameters, including oxygen consumption (VO_2_), heart rate, blood pressure, and respiratory exchange ratio (RER). An automated computerized ramp protocol was used to increase exercise intensity by 10 Watt/minute for women and 15 Watt/min for men, while participants were asked to maintain a constant velocity of 60 revolutions per minute. The highest average VO_2_ value recorded during an intensity interval (2/2.5 watts increment for women and men, respectively) was considered as the MAC value obtained from the procedure and that was used in further analyses.

### 2.4 MRI data acquisition

MRI scanning was performed at TASMC on a 3T Siemens system (MAGNETOM Prisma, Germany). Two high resolution anatomical sequences were implemented to improve further structural segmentation processes. T1-weighted images (voxel size = 1 × 1 × 1 mm) were acquired with a magnetization prepared 2 rapid acquisition gradient-echo protocol (MP2RAGE) with 176 contiguous slices using the following parameters: field of view (FOV) = 256 mm; repetition time (TR) = 5000 ms; echo time (TE) = 3.43 ms, inversion time 1 (TI1) = 803 ms, inversion time 2 (TI2) = 2500 ms, flip angle 1 (FA1) = 4°, flip angle 2 (FA2) = 5°. T2-weighted images (voxel size = 1 × 1 × 1 mm) were also acquired with 176 contiguous slices using the following parameters: field of view (FOV) = 256 mm; repetition time (TR) = 3200 ms; echo time (TE) = 413 ms. Diffusion weighted images (voxel size = 1.8 × 1.8 × 1.8 mm) were acquired along 20 directions using echo-planar-imaging (EPI) with 76 slices using the following parameters: field of view (FOV) = 240 mm; repetition time (TR) = 7200 ms; echo time (TE) = 55 ms, b value of 1000 s/mm^2^. Participants’ heads were stabilized with foam padding to minimize head movements. MP2RAGE images were reconstructed to create robust MP2RAGE T1w images without the strong background noise in air regions by implementing the regularization method proposed by 40.

### 2.5 MRI data analysis

#### 2.5.1 Structural MRI - hippocampal volume

Automatic subcortical volumetric segmentation was performed using FreeSurfer (v6.0 Martinos Center for Biomedical Imaging, Harvard-MIT, Boston, MA), which is documented and freely available for download (http://surfer.nmr.mgh.harvard.edu/). Both T1-weighted and T2-weighted images were used in this analysis. The technical details of these procedures were described previously (Fischl et al., 2002). Briefly, this fully automated process includes motion correction, removal of non-brain tissue, automated Talairach transformation, segmentation of the subcortical white matter and deep gray-matter volumetric structures (including hippocampus, amygdala, caudate, putamen, and ventricles), intensity normalization, and cortical reconstruction. This segmentation procedure assigns a neuroanatomical label to every voxel in the MR image volume. The method is based on probabilistic information estimated from a manually labeled training set. The Markov Random Field Theory is applied, where the probability of a label at a given voxel is computed not just in terms of the gray-scale intensities and prior probabilities at that voxel, but also as a function of the labels in a neighborhood around the voxel in question. This is particularly important for correct separation of the hippocampus and amygdala which have similar gray-scale values. For further data analysis the extracted bilateral hippocampal volumes were adjusted for estimated total intracranial volume (eTIV - provided by the automated pipeline of FreeSurfer), using SPSS by an analysis of covariance (ANCOVA) approach: adjusted volume = raw volume - β × (eTIV – mean eTIV), where β is the slope of a regression of the ROI volume on eTIV (Erickson et al., 2009; Raz et al., 2005; Raz et al., 2004). The resulting adjusted hippocampal volumes were then used as regions of interest in all subsequent analyses which were performed with statistical software.

#### 2.5.2 White matter lesions

White matter lesions volume was segmented and measured using the automatic pipeline of FreeSurfer, using both T1- and T2-weighted images as input. Importantly, a very strong correlation was recently demonstrated among cognitively-intact older adults between white matter hyper- and hypo-intense volume extracted from T2- and T1-weighted images, respectively, using the FreeSurfer pipeline ^41^. Furthermore, the WML burden based on T1 white matter hypointensities was found to exhibit stronger correlation with age, and β-amyloid concentration in the CSF, compared to WML volume based on T2 white matter hyperintensities. For further data analysis the extracted WML volume was adjusted for estimated total intracranial volume by an analysis of covariance (ANCOVA) approach identical to hippocampal volume.

#### 2.5.3 Diffusion MRI

Diffusion tensor imaging data were analyzed using the FMRIB Software Library (FSL) package. Preprocessing included correcting for eddy currents and head motion ^42^, and skull stripping that was performed using the Brain Extraction Tool (BET). Diffusion tensor models were then fitted using DTIfit, part of FSL Diffusion Toolbox. Next, we used the tract-based spatial statistics (TBSS) workflow ^43^ to non-linearly register all FA and MD maps into standard space, and the TBSS processed, non-skeletonized maps were then used in a ROI analysis approach using probabilistic atlases implemented in FSL. Hippocampal MD values were generated using a hippocampal mask created from the Harvard-Oxford atlas ^44^, with a 50% threshold. In addition, FA values of the fornix were extracted using a mask generated from the Jülich histological atlas ^45^, with a 50% threshold. The fornix is a primary afferent and efferent railway of information to and from the hippocampus ^46^, and structural deterioration of this tract has been shown to correlate with cognitive deficits ^15^. In order to minimize potential partial volume effects in hippocampal grey matter microstructural analysis ^5^, hippocampal masks were multiplied by a total grey matter segmentation mask thresholded at 70%, which was created with FSL’s FMRIB’s Automated Segmentation Tool (FAST) ^47^. One participant from the non-active group had corrupted diffusion data and was not included in all analyses incorporating DTI measures.

### 2.6 Cognitive assessment

Participants underwent memory assessment using two tests which evaluate distinct types of free delayed recall episodic memory. (I) In the Wechsler Logical Memory test (LM) ^48^ participants were asked to listen to two narrated stories and later, following 30 min, describe their plot details., (II) In the Rey Osterrieth Complex Figure test (ROCF) ^49^participants were asked to draw a copy of a complex abstract figure and later, after 20 min, draw it from their memory. The correct answers percentage of the two tests were then averaged to form a delayed-recall memory index which was used as the memory performance measure in the mediation analyses. Thirty-eight of the 50 participants (22 active/16 non-active) underwent the memory assessment and were included in all analyses implementing memory performance. In each analysis including memory assessment – age, sex, and education were controlled for.

### 2.7 Statistical analysis

Statistical analyses and visualizations were performed and constructed with IBM SPSS Statistics v24, and R 4.0.2. Since not all variables were normally distributed within each group, based on Kolmogorov-Smirnov test and histogram visual inspection, we tested differences between the active and non-active groups using permutation test. These procedures were conducted with 10,000 iterations, permuting each neurocognitive measure, and preserving the original group sizes. Then, the observed between-group means difference in each measure was compared to all between-group differences obtained from the permuted null distribution. P-values were determined as the probability of getting equal or greater between-group difference based on the null distribution. Between-group differences in continuous background characteristics (i.e., age, education, MAC) were evaluated with Wilcoxon-Mann-Whitney U test, while difference in the proportion between males and females were assessed using the Chi-squared test. To examine the relationships between MAC, hippocampal grey and white matter structural measures, WML burden, and memory performance we first generated adjusted values for each measure controlling for age, sex, and years of education, based on multiple regression unstandardized residuals. Then, we used permutation tests with 10,000 iterations to create a null distribution from which Pearson correlation coefficients were generated for each iteration. P-values were determined as the probability of getting equal or greater correlation coefficients compared to the observed correlation coefficient based on the permuted null distribution. Between-group tests’ and continuous correlations’ *p*-values were corrected for multiple tests using the False Discovery Rate (FDR) method ^50^. Since bilateral structural properties were highly correlated with each other (r>.75), all these measures were analyzed as total bilateral volume or average MD/FA. Then, based on the observed relationships between active lifestyle and MAC and all other variables (see *Results* section), a regression-based mediation analysis was conducted to test the mediating effect of white matter (WML and fornix microstructural integrity) and grey matter (hippocampal microstructural integrity) pathways on the relationship between active lifestyle/MAC and hippocampal macrostructure (i.e., volume). Next, a mediation analysis was used to examine biological paths mediating the relationship between active lifestyle/MAC and memory performance. Generally, in order to confirm a mediating effect of a proposed variable and its significance in the model, it needs to be demonstrated that while a significant association exists between the independent, dependent, and proposed mediating variables, the direct relationship between the independent and dependent variables is significantly reduced when the mediator (the indirect effect) is included in the model. The significance of the indirect mediating effects of WML, fornix FA, and hippocampal MD on the relationship between active lifestyle/MAC and hippocampal volume were evaluated using a 95% confidence interval of 5000 bootstrap samples, equivalent to a 2-tailed test. If the confidence interval limits excluded zero, the indirect mediating effect test was considered significant. Then, based on our directional hypothesis regarding the mediating pathway of active lifestyle/MAC on neurocognition, and the mediating effect observed between active lifestyle/MAC and hippocampal volume (see *Results* section), we used a 90% confidence interval, equivalent to a 1-tailed test, to examine the neuroprotective directional mediating pathways between active lifestyle/MAC and episodic memory. All measures were mean centered during these procedures. Mediation analyses were conducted using the macro-program PROCESS v3.5 implemented in SPSS ^51^. α≤.05 was set as the threshold for statistical significance in all analyses.

### 2.8 Data and code availability

Due to medical confidentiality and since participants did not consent to having their data stored in a public repository, the un-identified data (e.g., data spreadsheet) and code that support the findings of this study are available from the corresponding author, T.E., upon request. No formal data-sharing agreement is necessary.

## 3. Results

**Table 1** presents socio-demographic and cognitive characteristics of the whole sample and study sub-groups. The groups did not differ in any background parameter except for MAC, whereas expected the active group demonstrated significantly higher values.

**Table 1.**
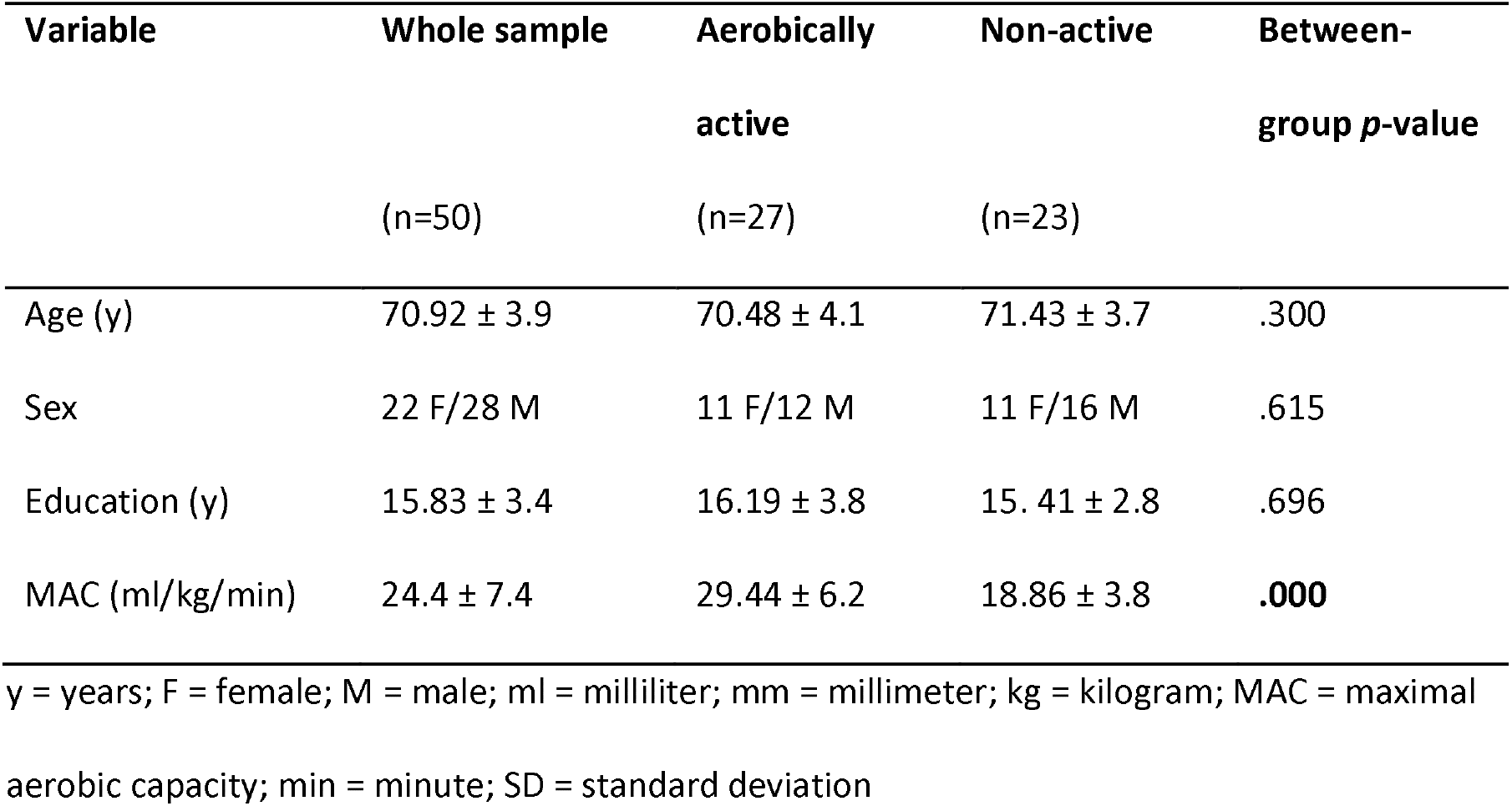
Socio-demographic and cognitive characteristics of participants.

### 3.1 Relationship between aerobically active lifestyle and neurobiological measures

The aerobically active group demonstrated increased hippocampal volume [difference=419.91 mm^3^, *p*=.023, *p*(FDR)=.027, **Figure 1a**], lower hippocampal MD [difference=−.08, *p*=.010, *p*(FDR)=.024, **Figure 1b**], higher fornix FA [difference=.02, *p*=.014, *p*(FDR)=.024, **Figure 1c**], and lower WML burden [difference=−495.22 mm^3^, *p*=.013, *p*(FDR)=.024, **Figure 1d**], compared to the non-active group.

**Figure 1.**
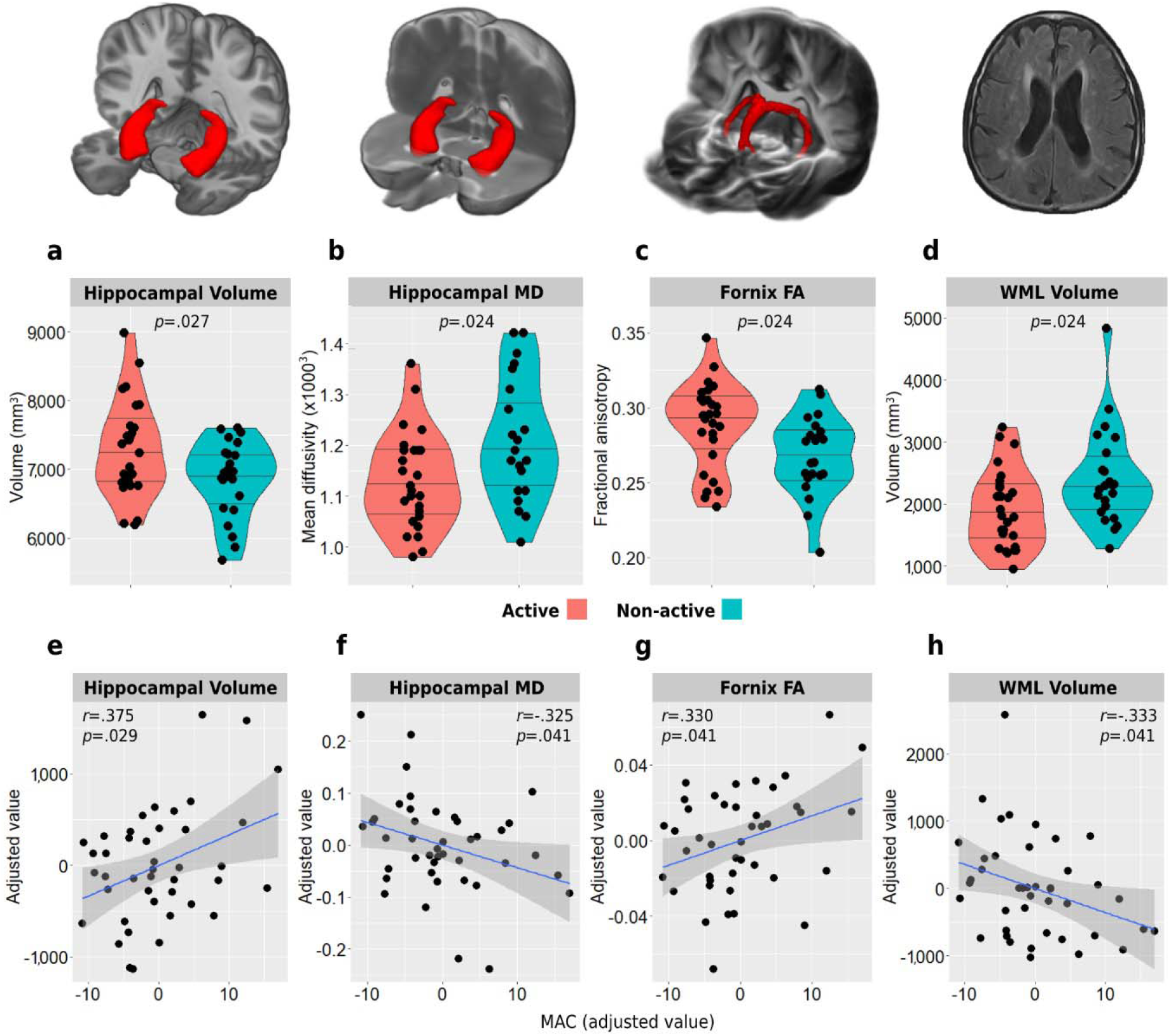
Between-group violin plots (a-d) and MAC correlations (e-h) with hippocampal grey and white matter structure, and WML burden. Correlations are adjusted for age, sex, and education. Horizontal lines in between-group plots represent 25%, 50%, and 75% quantiles. FA = fractional anisotropy; MAC = maximal aerobic capacity; MD = mean diffusivity; WML = white matter lesions

### 3.2 Relationship between MAC and neurobiological measures

A dose-response statistically significant moderate correlations were found between MAC and larger total hippocampal volume [*r*(40)=.375, *r^2^*=14%, *p*=.013, *p*(FDR)=.029, **Figure 1e**], lower hippocampal MD value [*r*(40)=−.325, *r^2^*=11%, *p*=.036, *p*(FDR)=.041, **Figure 1f**], increased fornix FA value [*r*(40)=.330, *r^2^*=11%, *p*=.033, *p*(FDR)=.041, **Figure 1g**], and lower WML volume [*r*(40)=−.333, *r^2^*=11%, *p*=.031, *p*(FDR)=.041, **Figure 1h**].

### 3.4 Relationship between aerobically active lifestyle and episodic memory

The aerobically active group was found to demonstrate statistically significant higher performance in the logical memory delayed recall test [difference=6.35, *p*=.006, *p*(FDR)=.024, **Figure 2a**] and the total episodic memory index [difference=8.41%, *p*=.023, *p*(FDR)=.027, **Figure 2c**] compared to the non-active group. While the active group also demonstrated a slight numerically higher performance in the ROCF delayed recall, this difference was not statistically significant [difference=1.51, *p*=.427, *p*(FDR)=.427, **Figure 2b**].

**Figure 2.**
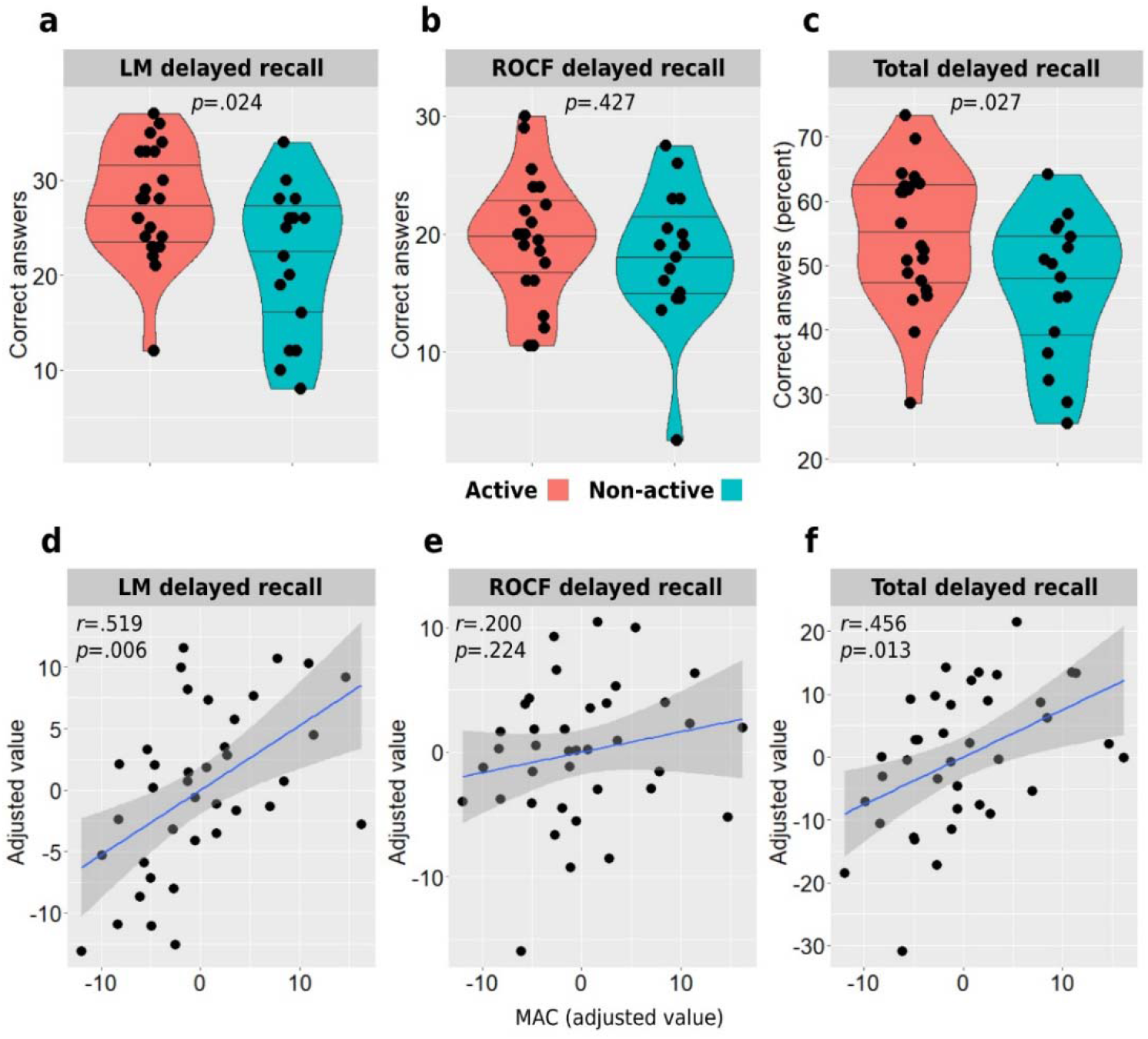
Between-group violin plots (a-c) and MAC correlations (d-f) with logical memory delayed recall, ROCF delayed recall, and total episodic memory index. Correlations are adjusted for age, sex, and education. Horizontal lines in between-group plots represent 25%, 50%, and 75% quantiles. LM = logical memory; ROCF = Rey Osterrieth Complex Figure

### 3.4 Relationship between MAC and episodic memory

MAC was also found to be significantly associated with logical memory delayed recall performance and the combined total episodic memory index, demonstrating statistically significant moderate correlations with these two measures [*r*(36)=.519, *r^2^*=27%, *p*=.001, *p*(FDR)=.006, and *r*(36)=.456, *r^2^* =21%, *p*=.004, *p*(FDR)=.013, respectively, **Figure 2d&f**]. In addition, MAC demonstrated weaker correlation with ROCF delayed recall performance, which did not reach statistical significance [*r*(36)=.200, *r^2^*=4%, *p*=.224, *p*(FDR)=.224, **Figure 2e**].

### 3.3 Examining mediating pathways between active lifestyle, and hippocampal volume and episodic memory

Between-group analyses revealed aerobically active lifestyle to be associated with all neurobiological measures, as well as episodic memory performance. In addition, hippocampal volume itself was also associated with all measures – hippocampal MD *r*(49)=−.476, *p*=.001; fornix FA *r*(49)=.557, *p*<.001; WML *r*(50)=−.415, *p*=.003; episodic memory index *r*(38)=.509, *p*=.001. Based on the aforementioned results, we first found lower hippocampal MD (partially standardized indirect effect=.30, SE=.15, 95% CI=[.05,.62]) to have a significant mediating effect on the relationship between active lifestyle and hippocampal macrostructure (i.e. volume) (**Figure 3a**). The total indirect effect accounted for ~47% of the total effect of aerobically active lifestyle on hippocampal volume, while the adjusted direct effect of MAC itself was statistically non-significant (*p*=.23). Next, since our neurobiological marker of cerebrovascular pathology is expressed as white matter damage, we examined whether a mediating pathway may exist between WML and fornix microstructure (FA value) in mediating the effect of active lifestyle on hippocampal volume. Using a serial mediation analysis we indeed found that fornix microstructure may constitute a significant link in the mediating pathway of WML on active lifestyle and hippocampal volume relationship (partially standardized indirect effect=.09, SE=.06, 95% CI=[.007,.24]) (**Figure 3b**). The total indirect effect accounted for ~70% of the total effect of MAC on hippocampal volume and was statistically significant (partially standardized indirect effect=.49, SE=.18, 95% CI=[.16,.86]), while the adjusted direct effect of active lifestyle itself was statistically insignificant (*p*=.46). We did not find a statistically significant mediating pathway between WML or fornix FA and hippocampal MD, suggesting for two independent mediating pathways between active lifestyle and hippocampal volume. Lastly, we used a serial mediation model and found that both pathways, i.e. WML-fornix FA-hippocampal volume (partially standardized indirect effect=.059, SE=.055, 90% CI=[.005,.17]) and hippocampal microstructure-hippocampal volume (partially standardized indirect effect=.16, SE=.10, 90% CI=[.03,.34]) significantly contributed to a mediating pathway between active lifestyle and episodic memory performance. In turn, these indirect pathways explained ~41% and ~49% of the total effect, respectively (**Figure 3c&d**).

**Figure 3.**
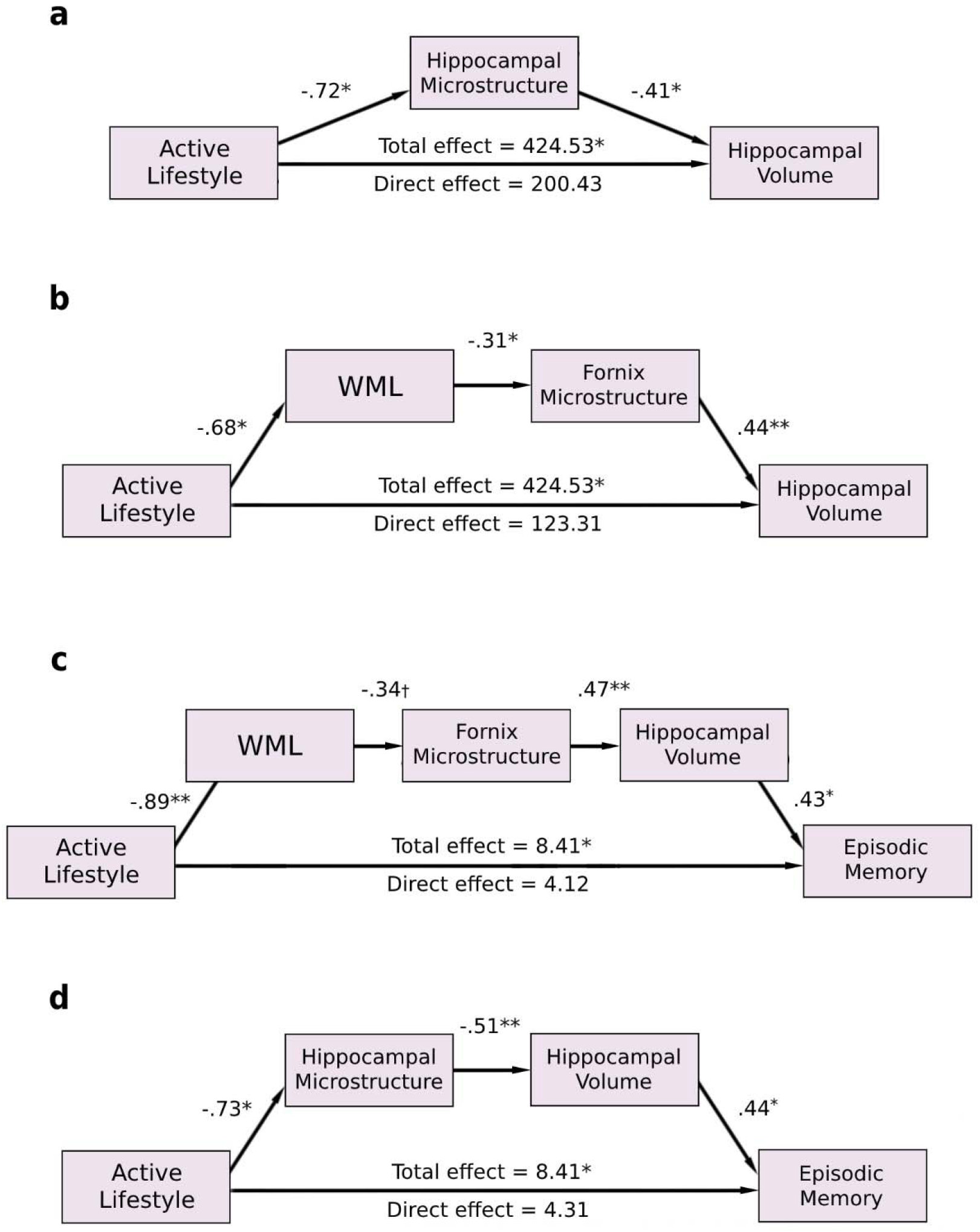
Mediation analysis revealing potential biological mechanistic pathways underlying the relationship between aerobically active lifestyle and hippocampal volume (a & b), and active lifestyle and episodic memory (c & d). Standardized coefficients are presented in the paths except in the total and direct effects. WML = white matter lesions; †*p*<0.1; * *p*<0.05; ** *p*<0.01

### 3.4 Examining mediating pathways between MAC, hippocampal volume and episodic memory

Based on the correlation analyses revealing MAC to be associated with all neurobiological measures, we found lower hippocampal MD (standardized indirect effect=.15, SE=.09, 95% CI=[.01,.35]) to have a mediating effect on the relationship between MAC and hippocampal macrostructure (i.e. volume) (**Figure 4a**). The indirect effect accounted for ~60% of the total effect of MAC on hippocampal volume, while the adjusted direct effect of MAC itself was statistically non-significant (*p*=.13). Next, since our neurobiological marker of cerebrovascular pathology is expressed as white matter damage, we examined whether a mediating pathway may exist between WML and fornix microstructure (FA value) in mediating the effect of MAC on hippocampal volume. Using a serial mediation analysis we indeed found that fornix microstructure may constitute a significant link in the mediating pathway of WML on MAC and hippocampal volume relationship (standardized indirect effect=.06, SE=.04, 95% CI=[.005,.15]) (**Figure 4b**). The total indirect effect accounted for ~60% of the total effect of MAC on hippocampal volume and was statistically significant (standardized effect=.22, SE=.10, 95% CI=[.03,.43]), while the adjusted direct effect of MAC itself was statistically insignificant (*p*=.28). We did not find a statistically significant mediating pathway between WML or fornix FA and hippocampal MD, suggesting for two independent mediating pathways between MAC and hippocampal volume. Lastly, we used a serial mediation model and found that both pathways, i.e. WML-fornix FA-hippocampal volume (standardized indirect effect=.04, SE=.03, 90% CI=[.003,.10]) and hippocampal microstructure-hippocampal volume (standardized indirect effect=.07, SE=.06, 90% CI=[.002,.19]) significantly contributed to a mediating pathway between MAC and episodic memory performance. In turn, these indirect pathways explained 41% and 42% of the total effect, respectively (**Figure 4c&d**).

**Figure 4.**
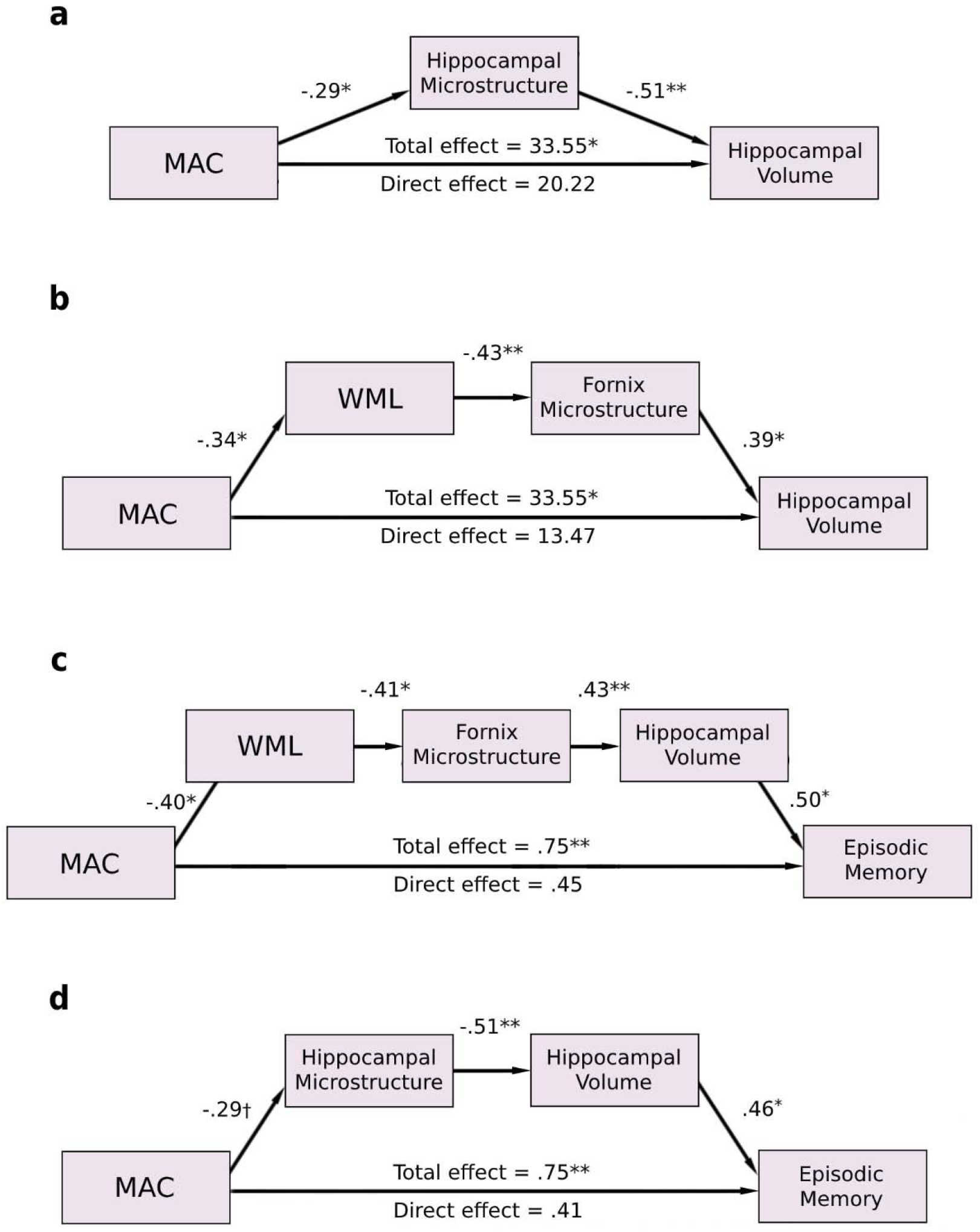
Mediation analysis revealing potential biological mechanistic pathways underlying the relationship between MAC and hippocampal volume (a & b), and MAC and episodic memory (c & d). Standardized coefficients are presented in the paths except in the total and direct effects. MAC = maximal aerobic capacity; WML = white matter lesions; †*p*<0.1; * *p*<0.05; ** *p*<0.01

## 4. Discussion

The present study aimed to investigate the relationship between aerobically active lifestyle and MAC, and hippocampal-related grey and white matter structure, as well as cerebrovascular pathology and memory performance in cognitively healthy older adults. We found aerobic exercise and MAC to have a neuroprotective relationship with hippocampal-related grey and white matter characteristics, i.e., hippocampal volume, and hippocampal and fornix microstructural integrity. In addition, aerobic activity and MAC were found to be associated with lower burden of WML volume, which adds evidence regarding the beneficial relationship between these lifestyle factors and reduced cerebrovascular pathology ^38^. These findings are in accordance with previous works that did find MAC- and aerobic exercise-related effect on the aged human hippocampus ^30–32,52^, and further strengthens the potential key role of MAC in the effect of aerobic activity on the brain. However, the biological mechanisms underlying these effects are not well understood. Thus, we also aimed to examine neurobiological pathways which may mediate the relationship between MAC and hippocampal volume using regression-based mediation analysis. We found two independent directional pathways which potentially mediate this relationship – a cerebrovascular-white matter path, and a hippocampal grey matter microstructural path. Furthermore, both of these pathways were found to mediate the relationship between MAC, hippocampal volume, and episodic memory.

Aging is associated with increased load of WML. The typical WML observed in older adults are presumed to be of vascular origin and constitute an expression of cerebral small vessels disease with neuropathological etiologies, such as hypoperfusion, increased blood-brain barrier permeability and deficits in vascular function ^16^. Although WML generally tend to be more localized in specific regions (e.g., periventricular and deep white matter), increased WML burden may also reflect a general pattern of increased pathological severity of the entire white matter microvasculature network. Furthermore, regions expressing WML have been found to be associated with not only pure vascular insult but with neural tissue changes including myelin and axonal lose ^53^, which may also indicate on a broader deterioration of other white matter areas integrity ^54,55^. This may include hippocampal-related white matter tracts as well, as was found in the current study by the significant negative relationship observed between WML volume and the fornix microstructural integrity. The fornix is a main bidirectional railway of information from and to the hippocampus, providing it with part of its neurochemical input, which is critical for its function ^46,56^. Hence, structural deterioration of the fornix may precedes and mediates volume loss of the hippocampus, as previously been demonstrated ^14,57^. This in turn, may explain the directional pathway we found from cerebrovascular-related white matter damage to fornix microstructural integrity in mediating the relationship between MAC and hippocampal volume. Our findings are also in accordance with previous works that found a positive relationship between MAC or aerobic exercise and hippocampal cerebrovascular integrity. In a study examining the effect of a 3-month aerobic exercise intervention in older adults, increased hippocampal perfusion was demonstrated to mediate the relationship between changes in aerobic fitness and hippocampal head volume and recognition memory performance ^31^. In animal studies neurogenesis and increased cell proliferation in the dentate gyrus of mice following an aerobic exercise period was found to correlate with increased blood volume and vessels density in this region, respectively ^58,59^. Since grey matter volume evaluated with MRI is a macroscale measure that is not restricted to cellular bodies alone, but also contains axonal components ^60^ and vascular structures ^61^, it is plausible that the mediating effect of healthier cerebrovasculature on hippocampal volume could also be expressed by increased density of hippocampal vasculature as been demonstrated in animals. Furthermore, decreased hippocampal angiogenesis but not neurogenesis has been found to be associated with decreased plasticity and aging in humans ^62^, further emphasizing the vital role of cerebrovascular integrity in the aged hippocampus. A contradictory finding comes from ^63^ who did not find a vascular explanation to the anterior hippocampal volume increase observed following a 6-week aerobic intervention in human adults using a multi-modal MRI approach. However, since the study sample examined by ^63^ was composed of much younger adults (mean age ~34) the results should be generalized to aged individuals with care.

While the hippocampus has been long recognized as a highly vulnerable brain area to impairments in blood supply and oxygen delivery ^64^, aerobic exercise have been associated not only with vascular but also oxygen-dependent energy production metabolic adaptations in the animal brain in general, and specifically in the hippocampus, including the promotion of mitochondrial function and biogenesis ^65,66^. Furthermore, MAC itself is a measure of aerobic metabolism, and higher MAC represents higher metabolic rate of the body’s aerobic energy production system. In addition, aerobic exercise has been found to promote increases in brain glycogen content in rats, including in the hippocampus ^67^. Glycogen is a metabolic reservoir of glucose in different tissues, including the brain, in which it is primarily located in astrocytes and was found to play key role in supporting brain activity and long-term memory ^68–70^. These features of metabolic correlates of aerobic exercise identified in animal brains suggest molecular mechanisms that could underlie the relationship between MAC and grey matter microstructural integrity. This in turn may (at least partially) explain the findings observed in the current study and also in ^32^ where changes in hippocampal MD were demonstrated to potentially mediate the relationship between improvement in aerobic fitness and increased hippocampal volume. This is further supported by the adverse effects aging may have on hippocampal mitochondrial physiology ^71^, which has been recognized to play a vital role in membrane breakdown and cellular integrity deterioration ^72^. Decreased membrane density, expansion of extracellular space, and cellular necrosis were all demonstrated to be associated with increased grey matter MD ^4,73^, further emphasizing its clinical significance in assessing grey matter structural integrity. Aerobic exercise has been also demonstrated to up-regulate neurotrophic factors secretion in the hippocampus, including brain-derived neurotrophic factor (BDNF) ^74,75^ and insulin-like growth factor-1 (IGF-1) which contribute to adult hippocampal neurogenesis and cell proliferation ^76,77^. In human older adults, increased BDNF levels following a year of aerobic intervention were found to be associated with increased bilateral anterior hippocampal volume, although its association with changes in aerobic fitness were not reported ^30^. Hence, these findings suggest that the relationship between MAC and hippocampal volume observed in humans may be partially driven by more preserved cellular density and integrity.

The present study suggests that the relationship between MAC and episodic memory may be mediated by parallel neurobiological pathways, including cerebrovascular, white matter, and grey matter mechanisms, and that it is not restricted to a single biological underpinning. These findings add evidence to the potential multidimensional effect that cardiorespiratory function, MAC and aerobic exercise may have on the brain and adds to our current limited knowledge on the neurobiological mediators of the neuroprotective effect of MAC in the aging human brain.

### 4.1 Study limitations

The main limitation of the current study is its cross-sectional nature. Although a well-accepted statistical method for investigating mediation effect revealed statistically significant directional mechanistic pathways, we cannot conclude any straight-forward causal relationship. Moreover, human studies are generally limited in their ability to directly investigate brain mechanisms due to the inaccessibility to directly measure neural tissue properties or induce lesions to test for causal dependency, thus correlative multimodal neuroimaging methodology followed by statistical mediation techniques may aid in shading more light on possible mechanistic pathways, which underlie neuroprotection in this case. Further research applying longitudinal and interventional methodology while testing for mechanistic mediation will add substantial value to these research questions.

## 5. Conclusions

Our study provides evidence for possible directional neurobiological pathways underlying the neuroprotective relationship observed between MAC and hippocampal volume, and its relation to episodic memory performance in cognitively healthy older adults. In addition, beyond the neurobiological value of our results, our findings provide further evidence to support and emphasize the potential significance of improved cardiorespiratory function and aerobically active lifestyle in promoting hippocampal-related neuroprotection and cognitive resilience in the aging human brain.

## Acknowledgements

We thank Prof. Dafna Ben bashat for her consultation during the research process. We also thank Dr. Moran Artzi for help with planning of the acquisition sequences, and Prof. Talma Hendler and Prof. Yuval Nir, for their thoughtful advice. In addition, we thank Dr. Avraham Man for help with the cardiopulmonary exercise assessments. The study has been funded from the grant of the Israel Science Foundation (ISF) provided to YL, and the financial support of the Sagol Family Foundation for Brain Research. The funding sources were not involved in the conduction of the research.

## References

1 Raz, N. et al. Regional brain changes in aging healthy adults: general trends, individual differences and modifiers. Cerebral cortex (New York, N.Y. : 1991) 15, 1676–1689, doi:10.1093/cercor/bhi044 (2005).

2 Allen, J. S., Bruss, J., Brown, C. K. & Damasio, H. Normal neuroanatomical variation due to age: the major lobes and a parcellation of the temporal region. Neurobiology of aging 26, 1245–1260; discussion 1279–1282, doi:10.1016/j.neurobiolaging.2005.05.023 (2005).

3 Salminen, L. E. et al. Regional age differences in gray matter diffusivity among healthy older adults. Brain imaging and behavior 10, 203–211, doi:10.1007/s11682-015-9383-7 (2016).

4 Van Camp, N. et al. A complementary diffusion tensor imaging (DTI)-histological study in a model of Huntington’s disease. Neurobiology of aging 33, 945–959, doi:https://doi.org/10.1016/j.neurobiolaging.2010.07.001 (2012).

5 Henf, J., Grothe, M. J., Brueggen, K., Teipel, S. & Dyrba, M. Mean diffusivity in cortical gray matter in Alzheimer’s disease: The importance of partial volume correction. NeuroImage. Clinical 17, 579–586, doi:10.1016/j.nicl.2017.10.005 (2018).

6 Nobis, L. et al. Hippocampal volume across age: Nomograms derived from over 19,700 people in UK Biobank. NeuroImage. Clinical 23, 101904, doi:10.1016/j.nicl.2019.101904 (2019).

7 Raz, N., Rodrigue, K. M., Head, D., Kennedy, K. M. & Acker, J. D. Differential aging of the medial temporal lobe: a study of a five-year change. Neurology 62, 433–438 (2004).

8 den Heijer, T. et al. Structural and diffusion MRI measures of the hippocampus and memory performance. NeuroImage 63, 1782–1789, doi:10.1016/j.neuroimage.2012.08.067 (2012).

9 Langnes, E. et al. Anterior and posterior hippocampus macro- and microstructure across the lifespan in relation to memory—A longitudinal study. Hippocampus 30, 678–692, doi:10.1002/hipo.23189 (2020).

10 Weston, P. S., Simpson, I. J., Ryan, N. S., Ourselin, S. & Fox, N. C. Diffusion imaging changes in grey matter in Alzheimer’s disease: a potential marker of early neurodegeneration. Alzheimer’s research & therapy 7, 47, doi:10.1186/s13195-015-0132-3 (2015).

11 Carlesimo, G. A., Cherubini, A., Caltagirone, C. & Spalletta, G. Hippocampal mean diffusivity and memory in healthy elderly individuals: a cross-sectional study. Neurology 74, 194–200, doi:10.1212/WNL.0b013e3181cb3e39 (2010).

12 Rogalski, E. et al. Age-related changes in parahippocampal white matter integrity: a diffusion tensor imaging study. Neuropsychologia 50, 1759–1765, doi:10.1016/j.neuropsychologia.2012.03.033 (2012).

13 Stoub, T. R. et al. Age-related changes in the mesial temporal lobe: the parahippocampal white matter region. Neurobiology of aging 33, 1168–1176, doi:10.1016/j.neurobiolaging.2011.02.010 (2012).

14 Pelletier, A. et al. Structural hippocampal network alterations during healthy aging: a multi-modal MRI study. Frontiers in aging neuroscience 5, 84, doi:10.3389/fnagi.2013.00084 (2013).

15 Fletcher, E. et al. Loss of fornix white matter volume as a predictor of cognitive impairment in cognitively normal elderly individuals. JAMA neurology 70, 1389–1395, doi:10.1001/jamaneurol.2013.3263 (2013).

16 Alber, J. et al. White matter hyperintensities in vascular contributions to cognitive impairment and dementia (VCID): Knowledge gaps and opportunities. Alzheimer’s & dementia (New York, N. Y.) 5, 107–117, doi:10.1016/j.trci.2019.02.001 (2019).

17 de Leeuw, F.-E. et al. Prevalence of cerebral white matter lesions in elderly people: a population based magnetic resonance imaging study. The Rotterdam Scan Study. Journal of Neurology, Neurosurgery & Psychiatry 70, 9–14, doi:10.1136/jnnp.70.1.9 (2001).

18 Habes, M. et al. White matter hyperintensities and imaging patterns of brain ageing in the general population. Brain : a journal of neurology 139, 1164–1179, doi:10.1093/brain/aww008 (2016).

19 Prins, N. D. et al. Cerebral White Matter Lesions and the Risk of Dementia. Archives of neurology 61, 1531–1534, doi:10.1001/archneur.61.10.1531 (2004).

20 Eckerström, C. et al. High white matter lesion load is associated with hippocampal atrophy in mild cognitive impairment. Dementia and geriatric cognitive disorders 31, 132–138, doi:10.1159/000323014 (2011).

21 Crane, D. et al. Gray matter blood flow and volume are reduced in association with white matter hyperintensity lesion burden: a cross-sectional MRI study. Frontiers in aging neuroscience 7, doi:10.3389/fnagi.2015.00131 (2015).

22 Raji, C. A. et al. White matter lesions and brain gray matter volume in cognitively normal elders. Neurobiology of aging 33, 834.e837–816, doi:10.1016/j.neurobiolaging.2011.08.010 (2012).

23 Nosheny, R. L. et al. Variables associated with hippocampal atrophy rate in normal aging and mild cognitive impairment. Neurobiology of aging 36, 273–282, doi:10.1016/j.neurobiolaging.2014.07.036 (2015).

24 Gattringer, T. et al. Vascular risk factors, white matter hyperintensities and hippocampal volume in normal elderly individuals. Dementia and geriatric cognitive disorders 33, 29–34, doi:10.1159/000336052 (2012).

25 Fiford, C. M. et al. White matter hyperintensities are associated with disproportionate progressive hippocampal atrophy. Hippocampus 27, 249–262, doi:10.1002/hipo.22690 (2017).

26 Rovio, S. et al. Leisure-time physical activity at midlife and the risk of dementia and Alzheimer’s disease. The Lancet. Neurology 4, 705–711, doi:10.1016/s1474-4422(05)70198-8 (2005).

27 Colcombe, S. & Kramer, A. F. Fitness effects on the cognitive function of older adults: a meta-analytic study. Psychological science 14, 125–130, doi:10.1111/1467-9280.t01-1-01430 (2003).

28 Colcombe, S. J. et al. Aerobic exercise training increases brain volume in aging humans. The journals of gerontology. Series A, Biological sciences and medical sciences 61, 1166–1170 (2006).

29 Firth, J. et al. Effect of aerobic exercise on hippocampal volume in humans: A systematic review and meta-analysis. NeuroImage 166, 230–238, doi:https://doi.org/10.1016/j.neuroimage.2017.11.007 (2018).

30 Erickson, K. I. et al. Exercise training increases size of hippocampus and improves memory. Proceedings of the National Academy of Sciences of the United States of America 108, 3017–3022, doi:10.1073/pnas.1015950108 (2011).

31 Maass, A. et al. Vascular hippocampal plasticity after aerobic exercise in older adults. Molecular psychiatry 20, 585–593, doi:10.1038/mp.2014.114 (2015).

32 Kleemeyer, M. M. et al. Changes in fitness are associated with changes in hippocampal microstructure and hippocampal volume among older adults. NeuroImage 131, 155–161, doi:10.1016/j.neuroimage.2015.11.026 (2016).

33 Niemann, C., Godde, B. & Voelcker–Rehage, C. Not only cardiovascular, but also coordinative exercise increases hippocampal volume in older adults. Frontiers in aging neuroscience 6, 170–170, doi:10.3389/fnagi.2014.00170 (2014).

34 Jonasson, L. S. et al. Aerobic Exercise Intervention, Cognitive Performance, and Brain Structure: Results from the Physical Influences on Brain in Aging (PHIBRA) Study. Frontiers in aging neuroscience 8, 336–336, doi:10.3389/fnagi.2016.00336 (2017).

35 Wilmore, J. H., Costill, D. L. & Kenney, W. L. Physiology of sport and exercise 6th edn, (Champaign, IL: Human Kinetics, 2008).

36 Barnes, J. N., Taylor, J. L., Kluck, B. N., Johnson, C. P. & Joyner, M. J. Cerebrovascular reactivity is associated with maximal aerobic capacity in healthy older adults. Journal of applied physiology (Bethesda, Md. : 1985) 114, 1383–1387, doi:10.1152/japplphysiol.01258.2012 (2013).

37 Bailey, D. M. et al. Elevated aerobic fitness sustained throughout the adult lifespan is associated with improved cerebral hemodynamics. Stroke 44, 3235–3238, doi:10.1161/strokeaha.113.002589 (2013).

38 Sexton, C. E. et al. A systematic review of MRI studies examining the relationship between physical fitness and activity and the white matter of the ageing brain. NeuroImage 131, 81–90, doi:10.1016/j.neuroimage.2015.09.071 (2016).

39 Voss, M. W., Vivar, C., Kramer, A. F. & van Praag, H. Bridging animal and human models of exercise-induced brain plasticity. Trends in cognitive sciences 17, 525–544, doi:10.1016/j.tics.2013.08.001 (2013).

40 O’Brien, K. R. et al. Robust T1-weighted structural brain imaging and morphometry at 7T using MP2RAGE. PloS one 9, e99676, doi:10.1371/journal.pone.0099676 (2014).

41 Wei, K. et al. White matter hypointensities and hyperintensities have equivalent correlations with age and CSF β-amyloid in the nondemented elderly. Brain and behavior 9, e01457, doi:10.1002/brb3.1457 (2019).

42 Andersson, J. L. R. & Sotiropoulos, S. N. An integrated approach to correction for off-resonance effects and subject movement in diffusion MR imaging. NeuroImage 125, 1063–1078, doi:10.1016/j.neuroimage.2015.10.019 (2016).

43 Smith, S. M. et al. Tract-based spatial statistics: voxelwise analysis of multi-subject diffusion data. NeuroImage 31, 1487–1505, doi:10.1016/j.neuroimage.2006.02.024 (2006).

44 Desikan, R. S. et al. An automated labeling system for subdividing the human cerebral cortex on MRI scans into gyral based regions of interest. NeuroImage 31, 968–980, doi:10.1016/j.neuroimage.2006.01.021 (2006).

45 Eickhoff, S. B., Heim, S., Zilles, K. & Amunts, K. Testing anatomically specified hypotheses in functional imaging using cytoarchitectonic maps. NeuroImage 32, 570–582, doi:10.1016/j.neuroimage.2006.04.204 (2006).

46 Cassel, J.-C., Duconseille, E., Jeltsch, H. & Will, B. THE FIMBRIA-FORNIX/CINGULAR BUNDLE PATHWAYS: A REVIEW OF NEUROCHEMICAL AND BEHAVIOURAL APPROACHES USING LESIONS AND TRANSPLANTATION TECHNIQUES. Progress in Neurobiology 51, 663–716, doi:https://doi.org/10.1016/S0301-0082(97)00009-9 (1997).

47 Zhang, Y., Brady, M. & Smith, S. Segmentation of brain MR images through a hidden Markov random field model and the expectation-maximization algorithm. IEEE transactions on medical imaging 20, 45–57, doi:10.1109/42.906424 (2001).

48 Kent, P. The Evolution of the Wechsler Memory Scale: A Selective Review. Applied Neuropsychology: Adult 20, 277–291, doi:10.1080/09084282.2012.689267 (2013).

49 Shin, M. S., Park, S. Y., Park, S. R., Seol, S. H. & Kwon, J. S. Clinical and empirical applications of the Rey-Osterrieth Complex Figure Test. Nature protocols 1, 892–899, doi:10.1038/nprot.2006.115 (2006).

50 Benjamini, Y. & Hochberg, Y. Controlling the False Discovery Rate: A Practical and Powerful Approach to Multiple Testing. Journal of the Royal Statistical Society. Series B (Methodological) 57, 289–300 (1995).

51 Hayes, A. F. & Little, T. D. Introduction to mediation, moderation, and conditional process analysis : a regression-based approach., (New York : The Guilford Press, 2018).

52 Erickson, K. I. et al. Aerobic fitness is associated with hippocampal volume in elderly humans. Hippocampus 19, 1030–1039, doi:10.1002/hipo.20547 (2009).

53 Gouw, A. A. et al. Heterogeneity of small vessel disease: a systematic review of MRI and histopathology correlations. Journal of neurology, neurosurgery, and psychiatry 82, 126–135, doi:10.1136/jnnp.2009.204685 (2011).

54 Chao, L. L. et al. Associations between white matter hyperintensities and β amyloid on integrity of projection, association, and limbic fiber tracts measured with diffusion tensor MRI. PloS one 8, e65175, doi:10.1371/journal.pone.0065175 (2013).

55 Leritz, E. C. et al. Associations between T1 white matter lesion volume and regional white matter microstructure in aging. Human brain mapping 35, 1085–1100, doi:10.1002/hbm.22236 (2014).

56 Hasselmo, M. E. Neuromodulation: acetylcholine and memory consolidation. Trends in cognitive sciences 3, 351–359, doi:https://doi.org/10.1016/S1364-6613(99)01365-0 (1999).

57 Metzler-Baddeley, C. et al. Fornix white matter glia damage causes hippocampal gray matter damage during age-dependent limbic decline. Scientific reports 9, 1060, doi:10.1038/s41598-018-37658-5 (2019).

58 Pereira, A. C. et al. An in vivo correlate of exercise-induced neurogenesis in the adult dentate gyrus. Proceedings of the National Academy of Sciences of the United States of America 104, 5638–5643, doi:10.1073/pnas.0611721104 (2007).

59 Van der Borght, K. et al. Physical exercise leads to rapid adaptations in hippocampal vasculature: temporal dynamics and relationship to cell proliferation and neurogenesis. Hippocampus 19, 928–936, doi:10.1002/hipo.20545 (2009).

60 Kassem, M. S. et al. Stress-induced grey matter loss determined by MRI is primarily due to loss of dendrites and their synapses. Molecular neurobiology 47, 645–661, doi:10.1007/s12035-012-8365-7 (2013).

61 Marinković, S., Milisavljević, M. & Puskas, L. Microvascular anatomy of the hippocampal formation. Surgical neurology 37, 339–349, doi:10.1016/0090-3019(92)90001-4 (1992).

62 Boldrini, M. et al. Human Hippocampal Neurogenesis Persists throughout Aging. Cell Stem Cell 22, 589–599.e585, doi:https://doi.org/10.1016/j.stem.2018.03.015 (2018).

63 Thomas, A. G. et al. Multi-modal characterization of rapid anterior hippocampal volume increase associated with aerobic exercise. NeuroImage 131, 162–170, doi:10.1016/j.neuroimage.2015.10.090 (2016).

64 Schmidt–Kastner, R. & Freund, T. F. Selective vulnerability of the hippocampus in brain ischemia. Neuroscience 40, 599–636, doi:https://doi.org/10.1016/0306-4522(91)90001-5 (1991).

65 Marques–Aleixo, I., Oliveira, P. J., Moreira, P. I., Magalhães, J. & Ascensão, A. Physical exercise as a possible strategy for brain protection: evidence from mitochondrial-mediated mechanisms. Prog Neurobiol 99, 149–162, doi:10.1016/j.pneurobio.2012.08.002 (2012).

66 Steiner, J. L., Murphy, E. A., McClellan, J. L., Carmichael, M. D. & Davis, J. M. Exercise training increases mitochondrial biogenesis in the brain. Journal of applied physiology (Bethesda, Md. : 1985) 111, 1066–1071, doi:10.1152/japplphysiol.00343.2011 (2011).

67 Matsui, T. et al. Brain glycogen supercompensation following exhaustive exercise. The Journal of physiology 590, 607–616, doi:10.1113/jphysiol.2011.217919 (2012).

68 Dienel, G. A. & Cruz, N. F. Contributions of Glycogen to Astrocytic Energetics during Brain Activation. Metabolic brain disease 30, 281–298 (2015).

69 Dienel, G. A. The metabolic trinity, glucose-glycogen–lactate, links astrocytes and neurons in brain energetics, signaling, memory, and gene expression. Neuroscience letters 637, 18–25, doi:10.1016/j.neulet.2015.02.052 (2017).

70 Suzuki, A. et al. Astrocyte-neuron lactate transport is required for long-term memory formation. Cell 144, 810–823, doi:10.1016/j.cell.2011.02.018 (2011).

71 Navarro, A. et al. Hippocampal mitochondrial dysfunction in rat aging. American journal of physiology. Regulatory, integrative and comparative physiology 294, R501–509, doi:10.1152/ajpregu.00492.2007 (2008).

72 Osellame, L. D., Blacker, T. S. & Duchen, M. R. Cellular and molecular mechanisms of mitochondrial function. Best practice & research. Clinical endocrinology & metabolism 26, 711–723, doi:10.1016/j.beem.2012.05.003 (2012).

73 Douaud, G. et al. In vivo evidence for the selective subcortical degeneration in Huntington’s disease. NeuroImage 46, 958–966, doi:https://doi.org/10.1016/j.neuroimage.2009.03.044 (2009).

74 Wrann, C. D. et al. Exercise induces hippocampal BDNF through a PGC-1α/FNDC5 pathway. Cell metabolism 18, 649–659, doi:10.1016/j.cmet.2013.09.008 (2013).

75 Berchtold, N. C., Chinn, G., Chou, M., Kesslak, J. P. & Cotman, C. W. Exercise primes a molecular memory for brain-derived neurotrophic factor protein induction in the rat hippocampus. Neuroscience 133, 853–861, doi:10.1016/j.neuroscience.2005.03.026 (2005).

76 Trejo, J. L., Carro, E. & Torres–Aleman, I. Circulating insulin-like growth factor I mediates exercise-induced increases in the number of new neurons in the adult hippocampus. The Journal of neuroscience : the official journal of the Society for Neuroscience 21, 1628–1634, doi:10.1523/jneurosci.21-05-01628.2001 (2001).

77 Liu, P. Z. & Nusslock, R. Exercise-Mediated Neurogenesis in the Hippocampus via BDNF. Frontiers in neuroscience 12, 52, doi:10.3389/fnins.2018.00052 (2018).

